# CRISPR Artificial Splicing Factors

**DOI:** 10.1101/431064

**Authors:** Nathaniel Jillette, Albert Cheng

**Affiliations:** The Jackson Laboratory for Genomic Medicine, Farmington, CT 06032, USA; Department of Genetics and Genome Sciences, University of Connecticut Health Center, Farmington, CT 06030, USA; Institute for Systems Genomics, University of Connecticut Health Center, Farmington, CT 06030, USA

**Author notes:** Correspondence [Manuscript version: 1.0. Get current version and plasmids at http://CasFx.org].

## Abstract

We report here the engineering of CRISPR Artificial Splicing Factors (CASFx) based on an RNA-targeting CRISPR/Cas system. We showed that simultaneous exon inclusion and exclusion can be induced at distinct targets by differential positioning of CASFx. We also created inducible CASFx (iCASFx) using the FKBP-FRB chemical-inducible dimerization domain, allowing small molecule control of alternative splicing.

## Main

Splicing is a process in which segments of a pre-mRNA called introns are removed while segments called exons are joined together to form mature mRNA^1^. Alternative splicing is a phenomenon in which different exon segments are spliced together to form mature mRNA with different sequences, greatly expanding the protein repertoire by allowing different proteins to be coded by a single gene. The process of alternative splicing is deeply embedded in gene regulatory networks by controlling gene isoform expression of > 90% of human genes^2^. Given such prevalence, dysregulation of (alternative) splicing has been implicated in many diseases^3–5^. RNA-seq is a powerful method to “read” transcriptomes for changes in alternative splicing in different cell types, conditions and diseases^2,5,6^. However, Scalable tools for precisely and reversibly “writing” alterative splicing is lacking.

Fusion of RNA regulatory proteins to heterologous RNA binding domains, such as Pumilio/PUF, MS2 coat protein (MCP), PP7 coat protein (PCP), and λN, have allowed artificial modulation of RNA processes^7–12^. For example, tethering of serine-rich (RS) domains or Glycine-rich (Gly) domains by engineered PUF domains to exons induce their inclusion or exclusion, respectively^9^. However, these artificial RNA effectors require either protein engineering or insertion of artificial tags to target RNA, and they depend on short recognition sequences, thus affording only limited targeting flexibility or specificity.

The fields of genetics and epigenetics have greatly benefited by the explosion of technologies based on RNA-guided DNA-targeting CRISPR/Cas systems^13^. We and others have successfully implemented molecular tools for modifying genetic sequences or epigenetic states of target DNA loci^14–22^. The exciting prospect of using CRISPR to target RNA was first demonstrated by conversion of the most frequently used DNA-targeting SpCas9 to an RNA nuclease “RCas9” with an addition of a PAMmer, an oligo when bound to target RNA mimics the Protospacer Adjacent Motif (PAM) required for SpCas9 binding^16^. Targeting of RCas9 to repetitive sequences does not require PAMmer^23^, however repeat sequences constitute only a small proportion of all RNA cis-regulatory elements. Following the initial report of RCas9, other CRISPR/Cas9 systems were also found to bind to single-stranded RNA in vitro^24^, 25, but evidence for their *in vivo* RNA binding in mammalian cells is lacking. RNA-guided RNA nucleases from bacterial CRISPR systems has recently been discovered^26–28^. Their adaptation to mammalian cells has not only allowed programmable RNA degradation^26,28,29^ but has also unleashed great potential for engineering novel tools for RNA-guided regulation of endogenous RNAs. These RNA nucleases showed superior specificity compared to RNAi^28^, and are amenable for engineering to create new functions, e.g., RNA sequence editing^27^, live cell RNA imaging^29^, and diagnostics^30^. In particular, CasRx is the most recently identified type IV-D CRISPR-Cas ribonuclease isolated from *Ruminococcus flavefaciens XPD3002* with robust activity in degrading target RNAs matching designed guide RNA (gRNA) sequences^28^. Furthermore, dCasRx with nuclease domains mutated (R239A/H244A/R858A/H863A) can be programmed to bind splicing elements to inhibit exon splicing, potentially by blocking access of splicing machinery. Induction of exon inclusion, however, has yet to be demonstrated.

In contrast to exon exclusion that can be sufficiently induced by binding of dCasRx alone^28^, we decided to create CASFx to induce exon inclusion which likely requires activity of splicing activator proteins^31^. We chose the Exon 7 of *SMN2* (*SMN2*-E7) as our test exon as it has implications in diseases and its regulation is well-characterized^32^. We created CRISPR Artificial Splicing Factors (CASFx) by fusing dCasRx with RBFOX1 or RBM38 splicing factors which were successfully applied in aptamer tethering assays to activate exon inclusion when bound downstream of the target exon in splicing minigenes^7, 11^. We created RBFOX1N-dCasRx-C by replacing segments containing RNA recognition motif (RRM) of splicing factor RBFOX1 (residues 118–189) with dCasRx and tested its activity to induce inclusion of *SMN2*-E7 in an *SMN2* splicing minigene (Fig 1A). Four guide RNAs (gRNAs g*SMN2*-1 through g*SMN2*-4) were designed within the intron between *SMN2*-E7 and E8. When cells were transfected with pCI-SMN2 (containing the splicing minigene) and control GFP plasmid (pmaxGFP), *SMN2* minigene expressed predominantly exclusion isoform (Fig 1B, lane 1). When transfected with RBFOX1N-dCasRx-C and individual *SMN2* intronic gRNAs, inclusion isoform level increased (Fig 1B, lanes 11∼14, see upper bands). Introduction of pools of two, three or four gRNAs simultaneously, further increased levels of E7-included transcripts, as well as deceased the levels of E7-excluded transcripts, switching the splicing pattern to predominantly inclusion (Fig 1B, lanes 15∼16). *SMN2*-E7 activation is dependent on RBFOX1 effector because dCasRx alone did not result in activation (Fig 1B, lanes 2∼9). Activation is also dependent on binding of the RBFOX1N-dCasRx-C on the *SMN2* intron as control gRNAs (“C”) did not induce *SMN2*-E7 inclusion (Fig 2, lanes 2 and 10). Since dCasRx binding within exon was shown to induce exon exclusion in a previous study, we asked whether RBFOX1N-dCasRx-C can also induce exon exclusion when bound within exon. We designed an exonic (“EX”) gRNA at the middle at *SMN2-*E7 (Fig 2A). We also created two additional CASFx by fusing RBM38 to N- or C-terminus of dCasRx, resulting in RBM38-dCasRx and dCasRx-RBM38, respectively. The *SMN2* minigene expressed predominantly the exclusion isoform in HEK293T cells transfected with control pmaxGFP plasmid (Fig 2B, lane 1). When transfected with one of the CASFx and the pool of DN gRNAs, *SMN2* showed a switch to predominantly inclusion isoform (Fig 2B, lanes 6,9,12). Again, *SMN2*-E7 activation was dependent on the RBFOX1 or RBM38 effectors because dCasRx alone did not result in activation (Fig 2B, lane 3). The activation was also dependent on CASFx binding to the *SMN2* intron as control guide did not induce exon inclusion (Fig 2B, lanes 5,8,11). Binding of CASFx within exon 7 induced its exclusion independent of effector fusion since dCasRx alone also induced exon exclusion when bound at this position (Fig 2B, lanes 4,7,10,13), consistent with a previous report^28^. This demonstrates that exon inclusion or exclusion can be induced by the same CASFx (e.g., RBFOX1N-dCasRx-C) by designing gRNA targeting different locations on target transcripts.

**Fig 1.**
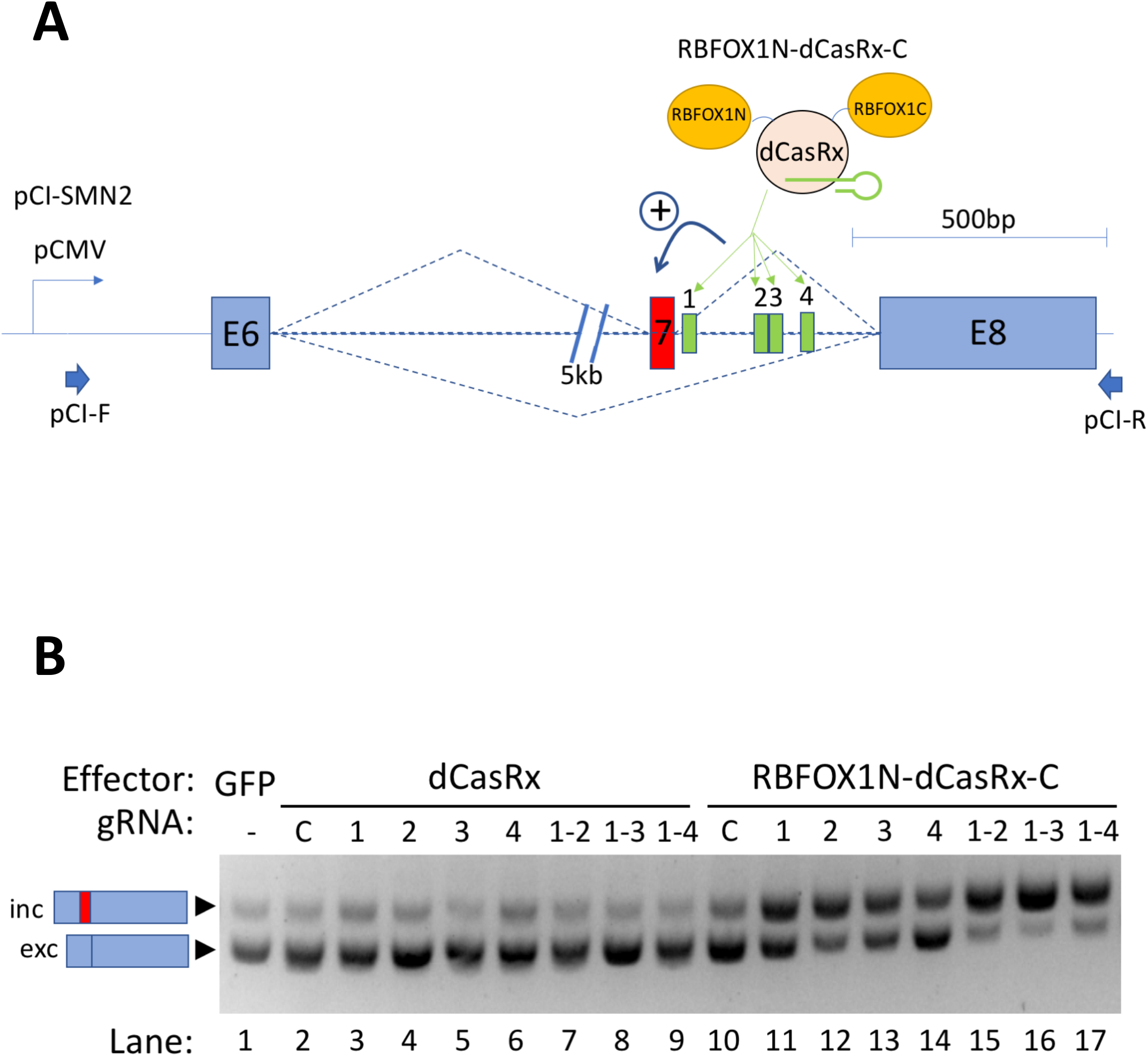
Exon inclusion induced by RBFOX1N-dCasRx-C. **(A)** Schematic of the artificial splicing factor RBFOX1N-dCasRx-C and *SMN2* minigene. The RNA binding domain of RBFOX1 was substituted by dCasRx to create an RNA-guided artificial splicing factor RBFOX1N-dCasRx-C that can be guided by guide RNAs (gRNA) to localize RBFOX1 splicing activity to target. The *SMN2* minigene on plasmid pCI-SMN2 contains exons 6 (E6) and 8 (E8) which are constitutively spliced, exon 7 (E7) that is alternatively spliced, and the intervening introns, driven by the CMV promoter (pCMV). Four designed target sites for the RBFOX1N-dCasRx-C are indicated by numbered boxes 1 through 4 within the intron between E7 and E8. pCI-F and pCIR indicate primers used for semi-quantitative RT-PCR assays. **(B)** Gel image of semi-quantitative splicing RT-PCR using primers pCI-F and pCI-R on *SMN2* minigene transcripts in cells co-transfected with control GFP plasmid (pmaxGFP), unfused dCasRx, or RBFOX1N-dCasRx-C, and the indicated guide RNAs (gRNAs). gRNA numbers correspond to those in Fig 1A with dash indicating the range of gRNAs used. “C” indicates a control gRNA without matching *SMN2* minigene sequence. Upper band and the lower band correspond to the exon 7-included and –excluded transcripts, respectively. **(C)** Column plots showing inc/exc ratio fold changes from quantitative RT-PCR (qRT-PCR) using primer pairs recognizing *SMN2* E7-included or –excluded isoforms.

**Fig 2.**
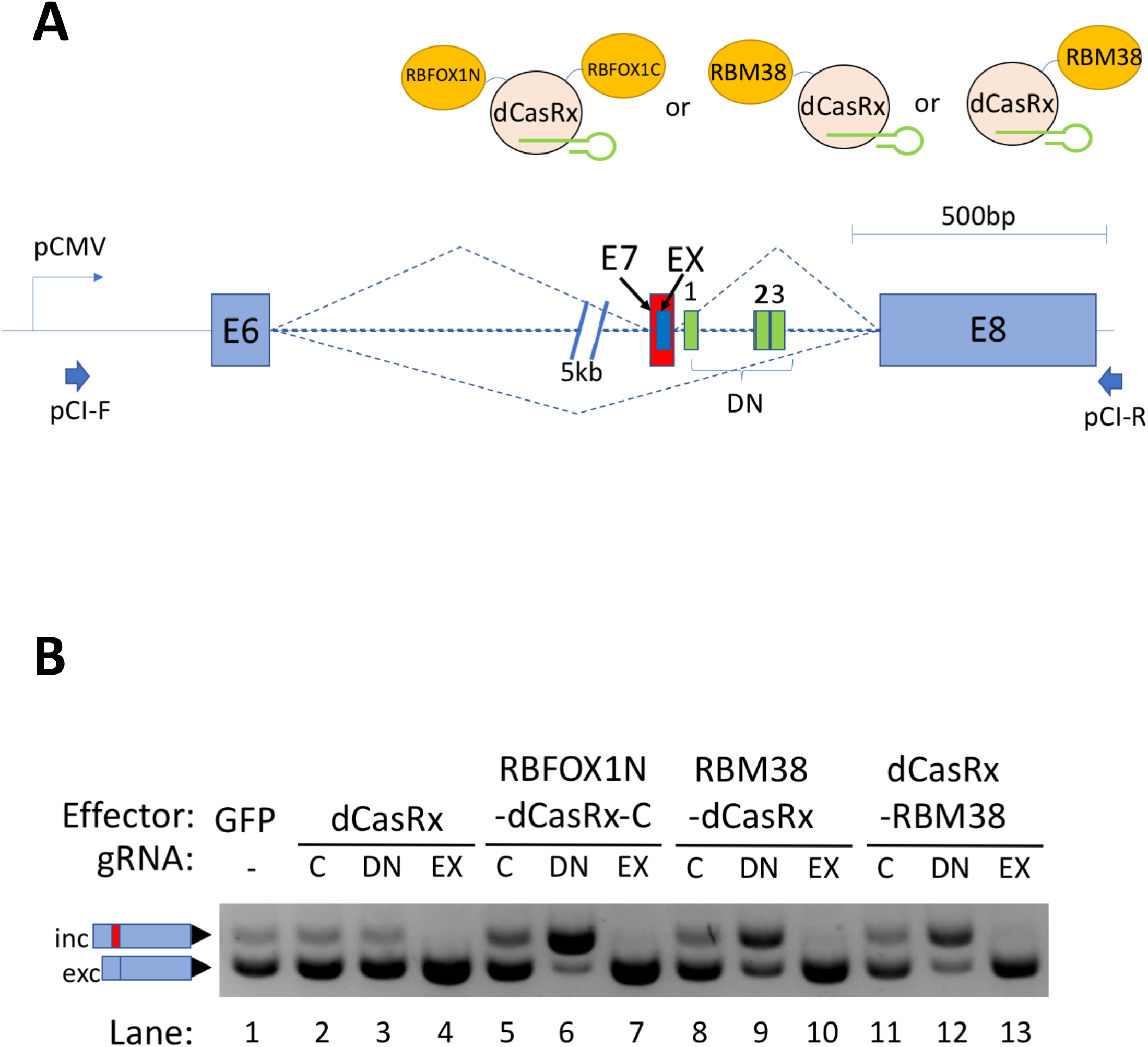
Activation and repression of exon by differential positioning of CASFx. **(A)** Schematic of the artificial splicing factors RBFOX1N-dCasRx-C, RBM38-dCasRx, dCasRx-RBM38 and *SMN2* minigene, as well as sets of three target sites (DN) downstream of E7 and one target site (EX) target within E7. **(B)** Gel image of semi-quantitative splicing RT-PCR using primers pCI-F and pCI-R on *SMN2* minigene transcripts in cells co-transfected with dCasRx, RBFOX1N-dCasRx-C, RBM38-dCasRx or dCasRx-RBM38, and the indicated gRNAs. “C” indicates a control gRNA without matching *SMN2* minigene sequence; “DN” indicates a pool of three gRNAs targeting downstream of E7; “EX” indicates a gRNA targeting within E7. Upper band and the lower band correspond to the exon 7-included and -excluded transcripts, respectively.

Next, we tested whether more than one splicing event can be modulated simultaneously and differently with CASFx (Fig 3A). We targeted RBFOX1N-dCasRx-C to the splice acceptor site of RG6 (RG6-SA) and DN locations at intron downstream of *SMN2*-E7 and observed simultaneous repression of RG6 cassette exon (RG6-CX) and activation of *SMN2*-E7 when both sets of gRNAs were transfected together with CASFx (Fig 3B, lane 4). Since CasRx is capable of processing gRNAs encoded in tandem (pre-gRNA) by cleaving 5’ of the direct repeats (DR) 28, we further tested whether the three SMN2-DN spacers and RG6-SA spacer could be encoded in one polycistronic pre-gRNA to achieve simultaneous modulation of the two splicing events. We first tested if the addition of a preprocessed DR to 3’ end of gRNA support CASFx function (Fig 3A, DR-SMN2–2-DR and DR-RG6-SA-DR). As predicted, these gRNAs remained active in inducing *SMN2*-E7 inclusion and RG6-CX exclusion (Fig 3B, lanes 5,6). More importantly, a polycistronic pre-gRNA (SMN2-DN-RG6-SA) harboring the three SMN2-DN spacers and the RG6-SA spacer induced simultaneous *SMN2*-E7 inclusion and RG6-CX exclusion when transfected with RBFOX1N-dCasRx-C (Fig 3B, lane 7), confirming the functionality of such polycistronic pre-gRNA architecture in inducing simultaneous and “bidirectional” splicing modulation of two different targets.

**Fig 3.**
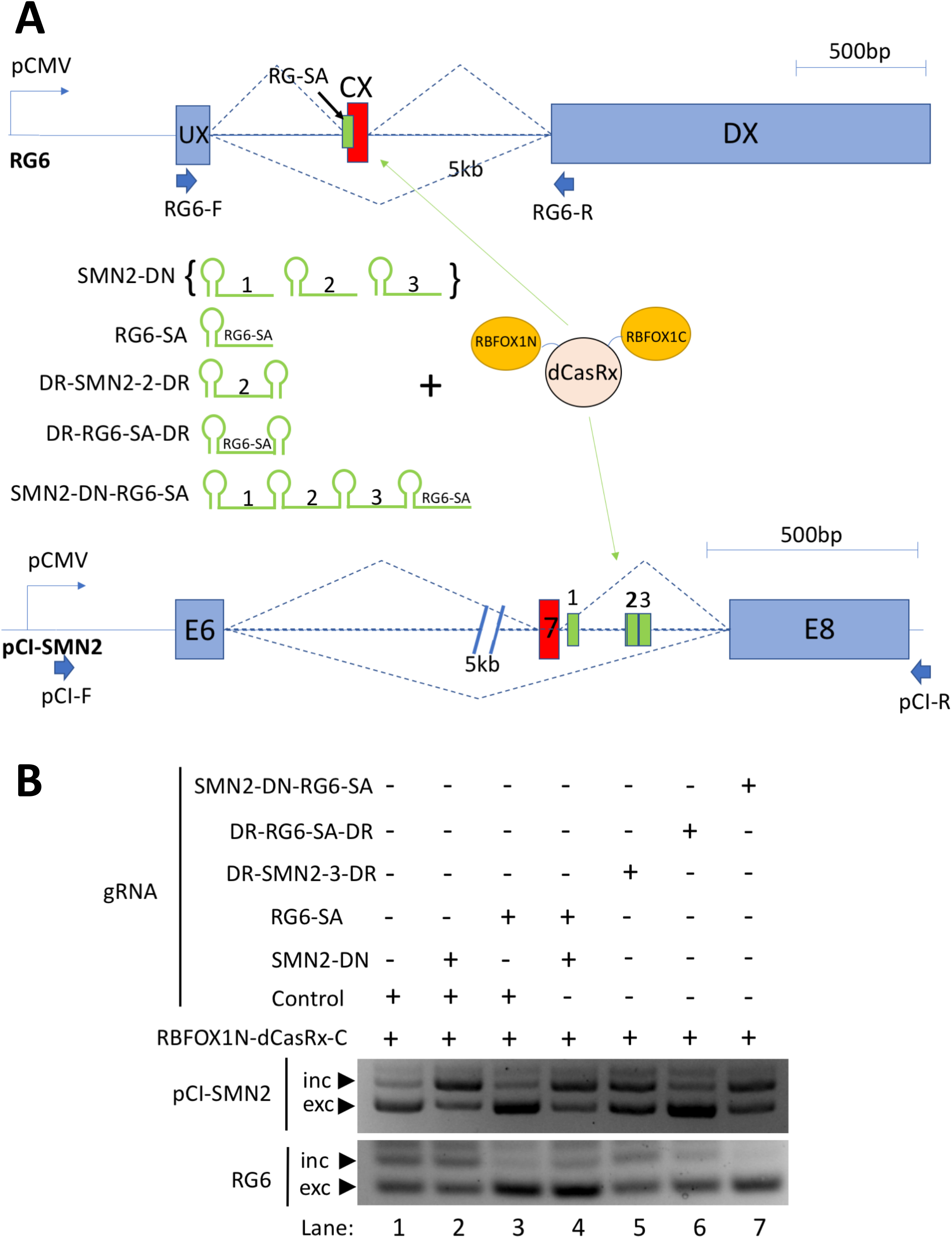
Simultaneous activation and repression of two independent exons by RBFOX1N-dCasRx-C directed by a polycistronic pre-gRNA. **(A)** Schematic of the artificial splicing factor RBFOX1N-dCasRx-C, various gRNA architectures, as well as the RG6 and *SMN2* minigenes. SMN2-DN gRNAs is a pool of three gRNAs, each expressed by a separate plasmid, targeting the corresponding numbered locations on the *SMN2* minigene. RG6-SA targets splice acceptor of RG6 cassette exon (CX). DR-SMN2–2-DR is SMN2 target 2 gRNA flanked by two direct repeats (DR). DR-RG6-SA-DR contains spacer against RG6-CX splice acceptor flanked by two DRs. SMN2-DN-RG6-SA is a polycistronic pre-gRNA with spacers targeting three DN sites on SMN2 downstream intron and RG6-CX splice acceptors intervened by DRs. **(B)** Gel image of semi-quantitative splicing RT-PCR of RG6 and *SMN2* minigene transcripts in cells cotransfected with the two minigene plasmids, RBFOX1N-dCasRx-C and the indicated gRNAs. Upper bands and the lower bands for the indicated transcripts correspond to the respective inclusion and exclusion isoforms.

Finally, we tested whether we could implement chemical control for CASFx. We created two-peptide inducible CRISPR Artificial Splicing Factors (iCASFx) by separating the RNA binding module (FKBP-dCasRx, or dCasRx-FKBP) and exon activation module (RBFOX1N-FRB-C, RBM38-FRB, or FRB-RBM38) into two peptides that can be induced to interact via the FKBPFRB domains^33^ in the presence of rapamycin (Fig 4A). Induction of *SMN2*-E7 inclusion was observed in cells cultured with rapamycin and transfected with iCASFx plus *SMN2*-DN-gRNA plasmids (Fig 4B, lanes 2,4,6,8,10,12) but not in cells cultured without rapamycin (Fig 4B, lanes 1,3,5,7,9,11).

**Fig 4.**
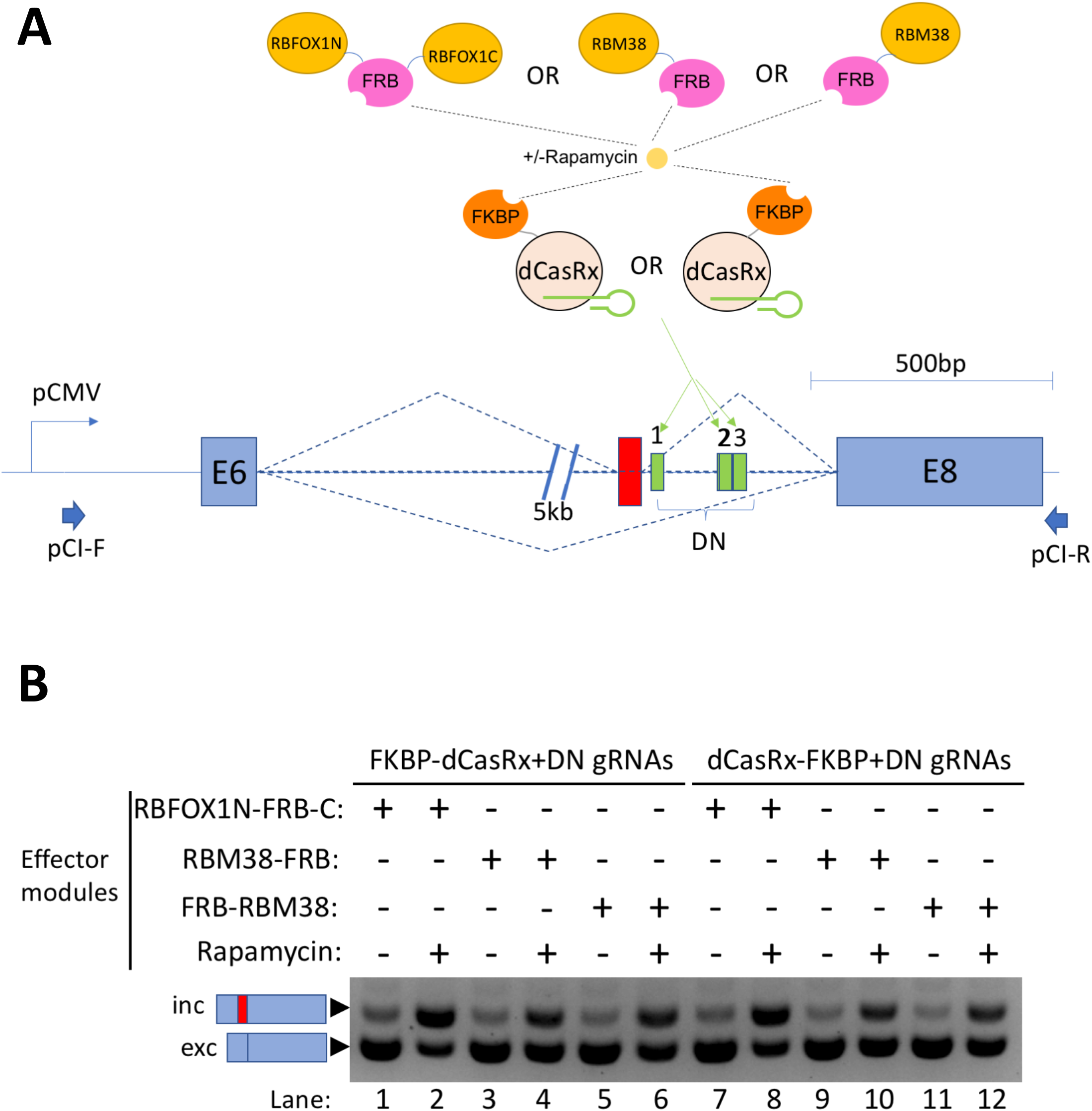
Chemical-inducible exon activation by two-peptide iCASFx. **(A)** Schematic of the two-peptide artificial splicing factors inducible by rapamycin. The RNA binding module (FKBP-dCasRx or dCasRx-FKBP) and effector module (RBFOX1N-FRB-C, RBM38-FRB, or FRBRBM38) containing the splicing activator domain are expressed separately as two peptides, fused to FKBP or FRB, respectively. FKBP and FRB can be induced to interact by rapamycin, bringing together the RNA binding module and the splicing activator module, and when guided by gRNAs, assemble at the target to activate exon inclusion. The *SMN2* minigene on plasmid pCI-SMN2 contains exons 6 (E6) and 8 (E8) which are constitutively spliced, exon 7 (E7) that is alternatively spliced, and the intervening introns, driven by the CMV promoter (pCMV). **(B)** Gel image of semi-quantitative splicing RT-PCR using primers pCI-F and pCI-R on *SMN2* minigene transcripts in cells co-transfected with the indicated constructs, and cultured with (“+”) or without (“-“) rapamycin. Upper band and the lower band correspond to the exon 7-included and -excluded transcripts, respectively.

In this study, we reported three RNA-guided splicing activators based on CRISPR/dCasRx. These CRISPR Artfiicial Splicing Factors (CASFx) can induce exon inclusion when targeted to bind at the downstream intron, and can induce exon exclusion when guided to bind within the target exon. We showed that simultaneous exon inclusion/exclusion can be achieved by a pool of gRNAs or a polycistronic pre-gRNA encoding spacers matching two target RNAs. Finally, we engineered inducible CRISPR Artificial Splicing Factors (iCASFx) that are inducible by small molecule rapamycin, potentially allowing spatiotemporal and tunable control of alternative splicing.

## Material & Methods

### Cloning

HEK293T cDNA was used as a source for PCR-amplification of coding sequences of splicing factors or other RNA binding proteins. Alternatively, gBlocks encoding human codon optimized versions of their coding sequences were ordered from IDT to serve as PCR template. The pXR002: EF1a-dCasRx-2A-EGFP ^28^ plasmid (Addgene #109050) served as PCR template for dCasRx coding sequence. The coding sequences of the CRISPR Artificial Splicing Factors (CASFx) were then cloned into pmax expression vector (Lonza) by a combination of fusion PCR, restriction-ligation cloning and Sequence- and Ligation-Independent Cloning (SLIC)^34^. gRNA expression cloning plasmids were generated by similar procedures using IDT oligonucleotides encoding CasRx gRNA direct repeat and PCR reaction using a ccdbCam selection cassette (Invitrogen) and a U6-containing plasmid as templates. Two BbsI restriction sites flanking the ccdbCam selection cassette serves as the restriction cloning sites for insertion of target-specific spacers. Target-specific spacer sequences were then cloned into the gRNA expression plasmids by annealed oligonucleotide ligation. Plasmid listing is included in the supplementary information. Plasmids and Genbank files will be available on Addgene. More supplementary data will also be on http://CasFx.org

### Cell culture and transfection

HEK293T cells were cultivated in Dulbecco’s modified Eagle’s medium (DMEM) (Sigma) with 10% fetal bovine serum (FBS)(Lonza), 4% Glutamax (Gibco), 1% Sodium Pyruvate (Gibco) and penicillin-streptomycin (Gibco). Incubator conditions were 37 °C and 5% CO2. For activation experiments, cells were seeded into 12-well plates at 100,000 cells per well the day before being transfected with 600ng (the “quota”) of plasmid DNA with 2.25uL Attractene tranfection reagent (Qiagen). 18 ng of each reporter minigene plasmid was transfected. The remaining quota was then divided equally among the effector and gRNA plasmids. In cases where there were two or more gRNA plasmids, the quota allocated for gRNA plasmids is further subdivided equally. For two-pepide effectors (i.e., the FKBP-FRB systems), the effector plasmid quota was divided equally between the plasmids encoding the individual peptides. Media was changed 24hr after transfection. 100nM (final concentration) of rapamycin was added during media change if applicable. Cells were harvested 48hr after transfection for RT-PCR analysis.

### RT-PCR

Cells were harvested for RNA extraction using RNeasy Plus Mini Kit (Qiagen). Equal amount of RNAs from one transfection experiment (either 700ng or 1000ng) were reverse-transcribed using High Capacity RNA-to-cDNA Kit (ThermoFisher). PCR was then performed using 2uL (out of 10uL) of cDNA using Phusion^®^ High-Fidelity DNA Polymerase (NEB) using minigene plasmid-specific primers for 25 cycles. PCR products were then analyzed on a 3% agarose gel.

## References

1. Wang, Z. & Burge, C.B. Splicing regulation: from a parts list of regulatory elements to an integrated splicing code. Rna 14, 802–813 (2008).

2. Wang, E.T. et al. Alternative isoform regulation in human tissue transcriptomes. Nature 456, 470 (2008).

3. Singh, R.K. & Cooper, T.A. Pre-mRNA splicing in disease and therapeutics. Trends in molecular medicine 18, 472–482 (2012).

4. Tazi, J., Bakkour, N. & Stamm, S. Alternative splicing and disease. Biochimica et Biophysica Acta (BBA)-Molecular Basis of Disease 1792, 14–26 (2009).

5. Park, E., Pan, Z., Zhang, Z., Lin, L. & Xing, Y. The expanding landscape of alternative splicing variation in human populations. The American Journal of Human Genetics 102, 11–26 (2018).

6. Shapiro, I.M. et al. An EMT–driven alternative splicing program occurs in human breast cancer and modulates cellular phenotype. PLoS genetics 7, e1002218 (2011).

7. Sun, S., Zhang, Z., Fregoso, O. & Krainer, A.R. Mechanisms of activation and repression by the alternative splicing factors RBFOX1/2. Rna 18, 274–283 (2012).

8. Choudhury, R., Tsai, Y.S., Dominguez, D., Wang, Y. & Wang, Z. Engineering RNA endonucleases with customized sequence specificities. Nature communications 3, 1147 (2012).

9. Wang, Y., Cheong, C.-G., Hall, T.M.T. & Wang, Z. Engineering splicing factors with designed specificities. Nature methods 6, 825 (2009).

10. Bos, T.J., Nussbacher, J.K., Aigner, S. & Yeo, G.W. in RNA Processing 61–88 (Springer, 2016).

11. Heinicke, L.A. et al. The RNA binding protein RBM38 (RNPC1) regulates splicing during late erythroid differentiation. PloS one 8, e78031 (2013).

12. Wang, X. et al. N 6-methyladenosine-dependent regulation of messenger RNA stability. Nature 505, 117 (2014).

13. Sander, J.D. & Joung, J.K. CRISPR-Cas systems for editing, regulating and targeting genomes. Nature biotechnology 32, 347 (2014).

14. Zalatan, J.G. et al. Engineering complex synthetic transcriptional programs with CRISPR RNA scaffolds. Cell 160, 339–350 (2015).

15. Hilton, I.B. et al. Epigenome editing by a CRISPR-Cas9-based acetyltransferase activates genes from promoters and enhancers. Nature biotechnology 33, 510 (2015).

16. O’connell, M.R. et al. Programmable RNA recognition and cleavage by CRISPR/Cas9. Nature 516, 263 (2014).

17. Qi, L.S. et al. Repurposing CRISPR as an RNA-guided platform for sequence-specific control of gene expression. Cell 152, 1173–1183 (2013).

18. Cheng, A.W. et al. Multiplexed activation of endogenous genes by CRISPR-on, an RNA-guided transcriptional activator system. Cell research 23, 1163 (2013).

19. Cheng, A.W. et al. Casilio: a versatile CRISPR-Cas9-Pumilio hybrid for gene regulation and genomic labeling. Cell research 26, 254 (2016).

20. Cong, L. et al. Multiplex genome engineering using CRISPR/Cas systems. Science, 1231143 (2013).

21. Morita, S. et al. Targeted DNA demethylation in vivo using dCas9–peptide repeat and scFv–TET1 catalytic domain fusions. Nature biotechnology 34, 1060 (2016).

22. Komor, A.C., Kim, Y.B., Packer, M.S., Zuris, J.A. & Liu, D.R. Programmable editing of a target base in genomic DNA without double-stranded DNA cleavage. Nature 533, 420 (2016).

23. Batra, R. et al. Elimination of toxic microsatellite repeat expansion RNA by RNA-targeting Cas9. Cell 170, 899–912. e810 (2017).

24. Strutt, S.C., Torrez, R.M., Kaya, E., Negrete, O.A. & Doudna, J.A. RNA-dependent RNA targeting by CRISPR-Cas9. Elife 7, e32724 (2018).

25. Rousseau, B.A., Hou, Z., Gramelspacher, M.J. & Zhang, Y. Programmable RNA cleavage and recognition by a natural CRISPR-Cas9 system from Neisseria meningitidis. Molecular cell 69, 906–914. e904 (2018).

26. Abudayyeh, O.O. et al. C2c2 is a single-component programmable RNA-guided RNA-targeting CRISPR effector. Science 353, aaf5573 (2016).

27. Cox, D.B. et al. RNA editing with CRISPR-Cas13. Science 358, 1019–1027 (2017).

28. Konermann, S. et al. Transcriptome engineering with RNA-targeting type VI-D CRISPR effectors. Cell 173, 665–676. e614 (2018).

29. Abudayyeh, O.O. et al. RNA targeting with CRISPR–Cas13. Nature 550, 280 (2017).

30. Gootenberg, J.S. et al. Nucleic acid detection with CRISPR-Cas13a/C2c2. Science, eaam9321 (2017).

31. Chen, M. & Manley, J.L. Mechanisms of alternative splicing regulation: insights from molecular and genomics approaches. Nature reviews Molecular cell biology 10, 741 (2009).

32. Cartegni, L. & Krainer, A.R. Disruption of an SF2/ASF-dependent exonic splicing enhancer in SMN2 causes spinal muscular atrophy in the absence of SMN1. Nature genetics 30, 377 (2002).

33. Luker, K.E. et al. Kinetics of regulated protein–protein interactions revealed with firefly luciferase complementation imaging in cells and living animals. Proceedings of the National Academy of Sciences 101, 12288–12293 (2004).

34. Jeong, J.-Y. et al. One-step sequence-and ligation-independent cloning (SLIC): rapid and versatile cloning method for functional genomics studies. Applied and environmental microbiology, AEM. 00844–00812 (2012).

